# Mechanisms of signalling-memory governing progression through the eukaryotic cell cycle

**DOI:** 10.1101/2020.08.20.259671

**Authors:** Béla Novák, John J Tyson

## Abstract

As cells pass through each replication-division cycle, they must be able to postpone further progression if they detect any threats to genome integrity, such as DNA damage or misaligned chromosomes. Once a ‘decision’ is made to proceed, the cell unequivocally enters into a qualitatively different biochemical state, which makes the transitions from one cell cycle phase to the next switch-like and irreversible. Each transition is governed by a unique signalling network; nonetheless, they share a common characteristic of bistable behaviour, a hallmark of molecular memory devices. Comparing the cell cycle signalling mechanisms acting at the Restriction Point, G1/S, G2/M and meta-to-anaphase transitions, we deduce a generic network motif of coupled positive and negative feedback loops underlying each transition.

## Introduction

The perpetuation of life depends on precise replication and segregation of chromosomes as cells pass through the cell division cycle. In eukaryotes, DNA replication and chromosome segregation are carried out in non-overlapping phases (S and M, respectively) with gap phases in between (G1-S-G2-M). It is essential that both chromosomal events happen once and only once between two successive cell divisions and that replication of the genome always precedes its segregation [1].

To guarantee the strict alternation of chromosome replication and segregation, cell cycle progression is controlled by **decision points** where a cell assesses its progress so far and commits itself to transition into the next stage of the cycle. There are four characteristic decision points and corresponding **cell cycle transitions**. Two of the decisions/transitions are associated with initiation of DNA replication (**G1/S**) and entering into mitosis (**G2/M**). The commitment to proliferate is made at the Restriction Point (**RP**), while chromosome segregation is determined at the meta-to-anaphase (**M/A**) transition of mitosis. Successful completion of the cell cycle requires that progression through these transitions is unidirectional: RP → G1/S → G2/M → M/A → … In a sense, the cell ‘remembers’ which stages it has already completed and moves to the next phase in the sequence. This unidirectionality—this memory—is guaranteed by the irreversible nature of the transitions, which is a property of the complex molecular signalling mechanisms operating at each transition. If any crucial event of the cell division cycle is compromised, the control machinery must halt further progression through the cycle and ‘remember’ where the cell is in the sequence, so that the cycle can resume exactly where it left off. If this memory is lost at any point of the cycle, earlier stages of the cycle may be repeated, leading to increase of ploidy or random segregation of chromosomes. The mechanisms responsible for the maintenance of cell-cycle memory are, thus, of fundamental importance to the integrity of the genome and the propagation of life.

Much is known about the molecular machinery controlling progression through the cell division cycle. Cell cycle transitions are triggered by ‘activator’ molecules whose levels fluctuate throughout the cell cycle (Figure 1A). The first three transitions are driven by different cyclin-dependent protein kinase complexes (Cyclin:Cdk, in our notation). In higher eukaryotes, CycE:Cdk2 acts at the RP, CycA:Cdk2 appears at G1/S, while CycB:Cdk1 is activated at the G2/M transition. The M/A transition is triggered by activation of a ubiquitin ligase, the Anaphase Promoting Complex/Cyclosome (APC/C), in partnership with Cdc20 (APC/C:Cdc20, in our notation). As the cell exits mitosis, powerful protein phosphatases reverse the phosphorylations carried out by the cyclin-dependent kinases and reset the control system to its starting point in early G1.

**Figure 1:**
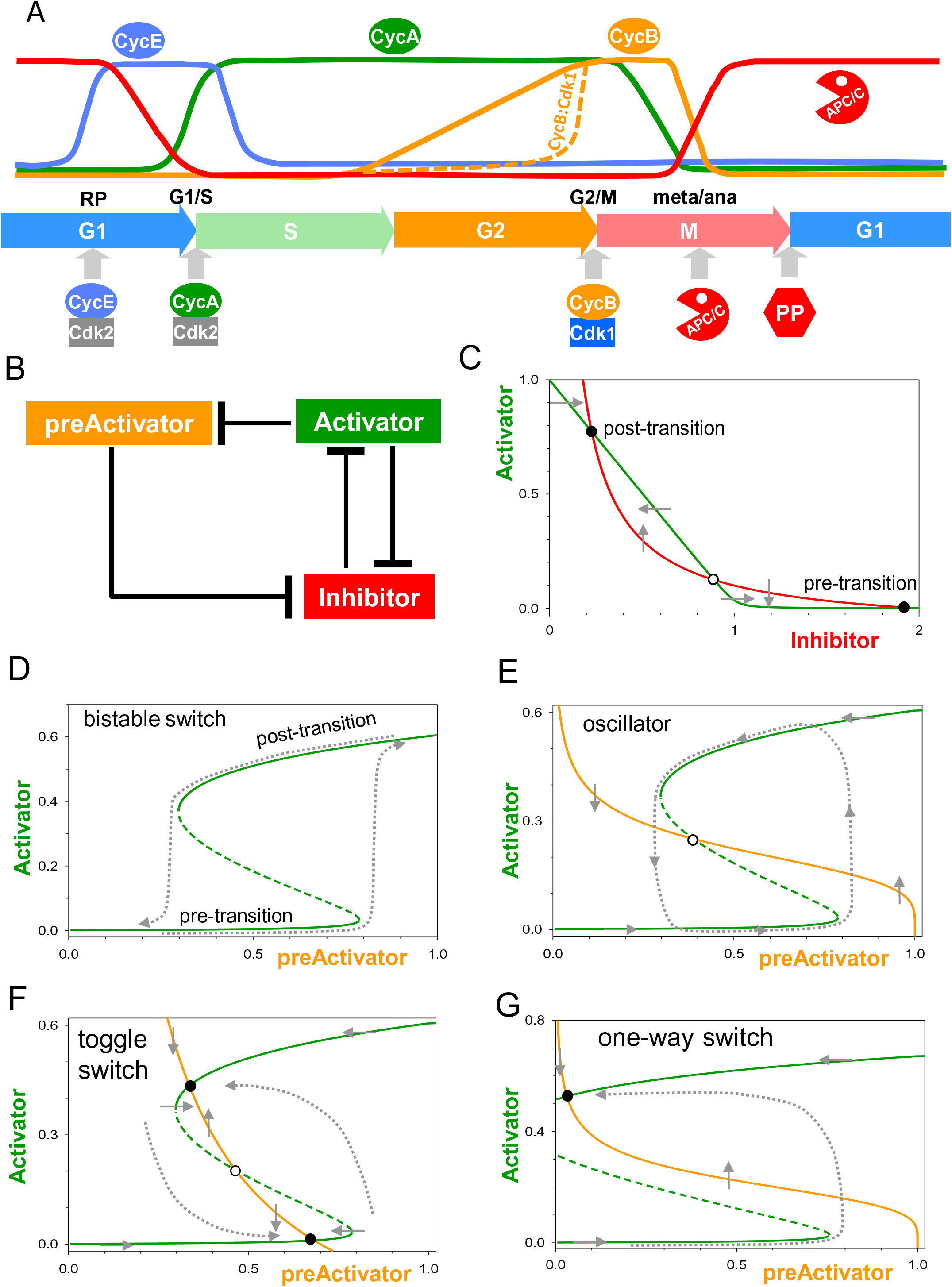
A generic network motif underlying irreversible cell cycle transitions. (A) Temporal oscillations of cell cycle activators in human cells (based on [1]). (B) Influence diagram of the network motif with a double-negative feedback loop embedded within a three-component negative feedback loop. The blunt connectors represent inhibitory signals. (C) Balance curves of Inhibitor (I) and Activator (A) involved in a double-negative feedback loop. Along a balance curve, the production and consumption rates of a component are equal to each other (‘balanced’); therefore, the concentration of the component is unchanging in time. For example, along the Inhibitor balance curve (red), dI/dt=0 so I(t) is not changing, although A(t) may be increasing or decreasing, as indicated by the grey vertical arrows. Similarly, along the Activator balance curve (green), A(t) is momentarily constant, but I(t) may be changing (horizontal grey arrows). The Inhibitor balance curve is a simple hyperbola, because A degrades I. The activator balance curve is a ‘hockey stick’, because I is a stoichiometric inhibitor of A. At each of the three intersections of the balance curves both dI/dt=0 and dA/dt=0, i.e., the control system is at a steady state. Two of the steady states are stable (●), and the intermediate steady state is unstable (○). (D) Signal-response curve (SRC) for the double-negative feedback loop. In general, a SRC depicts the effect of increasing a signal (an experimentally adjustable stimulus) on the long-term response in some characteristic feature of the control system. In this case, we are thinking of the concentration of the previous Activator (pA) as the signal, and the number and nature of the steady states of the next Activator (A) as the response. Solid lines denote stable steady states of A, and dashed lines, unstable steady states. When pA is small, 0 < pA < 0.25, steady-state activator level is very small; *i*.*e*., the lower branch of the curve represents the pre-transition state. When pA is large, pA > 0.8, activator level is large, which represents the post-transition state. The system is bistable for 0.25 < pA < 0.8, and it makes an abrupt transition from the pre-transition state to the post-transition state when pA passes the threshold ≈ 0.8, as indicated by the right-most dotted curve. When pA drops in the post-transition state, A level stays high until pA drops below the threshold ≈ 0.25, indicated by the left-most dotted curve. (E-G) The dynamics of the generic network motif is characterized by the balance curves for preActivator and Activator. The Activator balance curve (green) is the locus of points where dA/dt=0 for fixed values of pA, which is just the SRC plotted in panel D. The preActivator balance curve (orange) is a decreasing function of A because A is an inhibitor of pA. The grey arrows, as before, indicate the directions of changes of A(t) and pA(t) in time. The two balance curves can intersect three different ways, all of them being relevant for cell cycle transitions. If the SRCs intersect on the middle branch of the bistable switch, the network exhibits limit cycle oscillations indicated by the dotted closed trajectory in panel E. If they intersect in three places (F), two stable steady states and one unstable, the system functions as a toggle switch. In this case, sufficiently large perturbations away from either stable steady state can drive the system to the other stable steady state, as indicated by the dotted trajectories. The third case (G) corresponds to very strong double-negative feedback, so that the bistable switch stays in the active state even if pA drops to zero, as indicated again by the dotted trajectory. This is the case of a fully irreversible (one-way) switch. See Supplementary data S1 for details of calculations.

The temporal oscillations of cell cycle activators (Figure 1A) suggest a general **organizing principle** for the dynamics of the transitions. Each cell cycle activator helps to activate the next one in the sequence, and, after the transition, the previous activator is down-regulated while the new activator becomes stabilized. Neighbouring pairs of cell cycle activators underlying cell cycle transitions interact by a **generic network motif** of coupled negative and double-negative feedback loops (Figure 1B). At each transition, the activator of the previous transition (pA) helps the activator of the current transition (A) to eliminate its inhibitor (I), and the subsequent rise of A drives the cell into the next stage of the cell cycle. Subsequently, A drives down pA, but the mechanism maintains itself in the high A state because the transition is based on a bistable switch generated by the double-negative feedback between A and I. In this context, ‘bistability’ refers to two alternative steady states with low and high A level corresponding to pre- and post-transition states (Figure 1C). The transition from low to high A state requires pA to exceed a threshold level; thereafter, the double-negative feedback between A and I stabilizes the high A state, even as the activity of pA drops (Figure 1D).

As we will show, the alternative steady states of a bistable switch are responsible for the irreversibility of each transition in the cell cycle, i.e., for maintaining a memory of each transition. The molecular mechanisms of these switches are based on the generic motif in Figure 1B. The motif consists of a bistable switch (A **⊣**I **⊣**A) embedded in a negative feedback loop (pA **⊣**I **⊣**A **⊣**pA). Of central importance to our story is recognition that this motif can generate three different scenarios: oscillation (Figure 1E), toggle switch (Figure 1F) and one-way switch (Figure 1G).

### G2/M transition

Entry into mitosis is controlled by a mechanism based on phosphorylation of the CycB:Cdk1 complex (Activator): CycB:Cdk1 is inhibited by phosphorylation of the Cdk1 subunit by Wee1 kinase (Inhibitor) and re-activated by dephosphorylation by Cdc25 phosphatase (co-Activator) (Figure 2A). Because active CycB:Cdk1 inhibits Wee1 (double-negative feedback) and activates Cdc25 (double-positive feedback), the transition is governed by a bistable switch between low and high activities of CycB:Cdk1. The pA at this transition is CycA:Cdk1/2, which is not subject to inhibitory Cdk-phosphorylation by Wee1 [2]. CycB:Cdk1 was the first cell cycle activator proposed to be regulated by a bistable switch [3,4]. As a consequence of bistability, more CycB is required to activate Cdk1 than is the amount needed to maintain Cdk1 active (Figure 2B), as was experimentally confirmed in *Xenopus* cell-free extracts [5,6]. The phosphatase opposing CycB:Cdk1 on Wee1 and Cdc25 plays a critical role in creating bistability and in determining the thresholds for CycB:Cdk1 activation and inactivation (Figure 2C).

**Figure 2:**
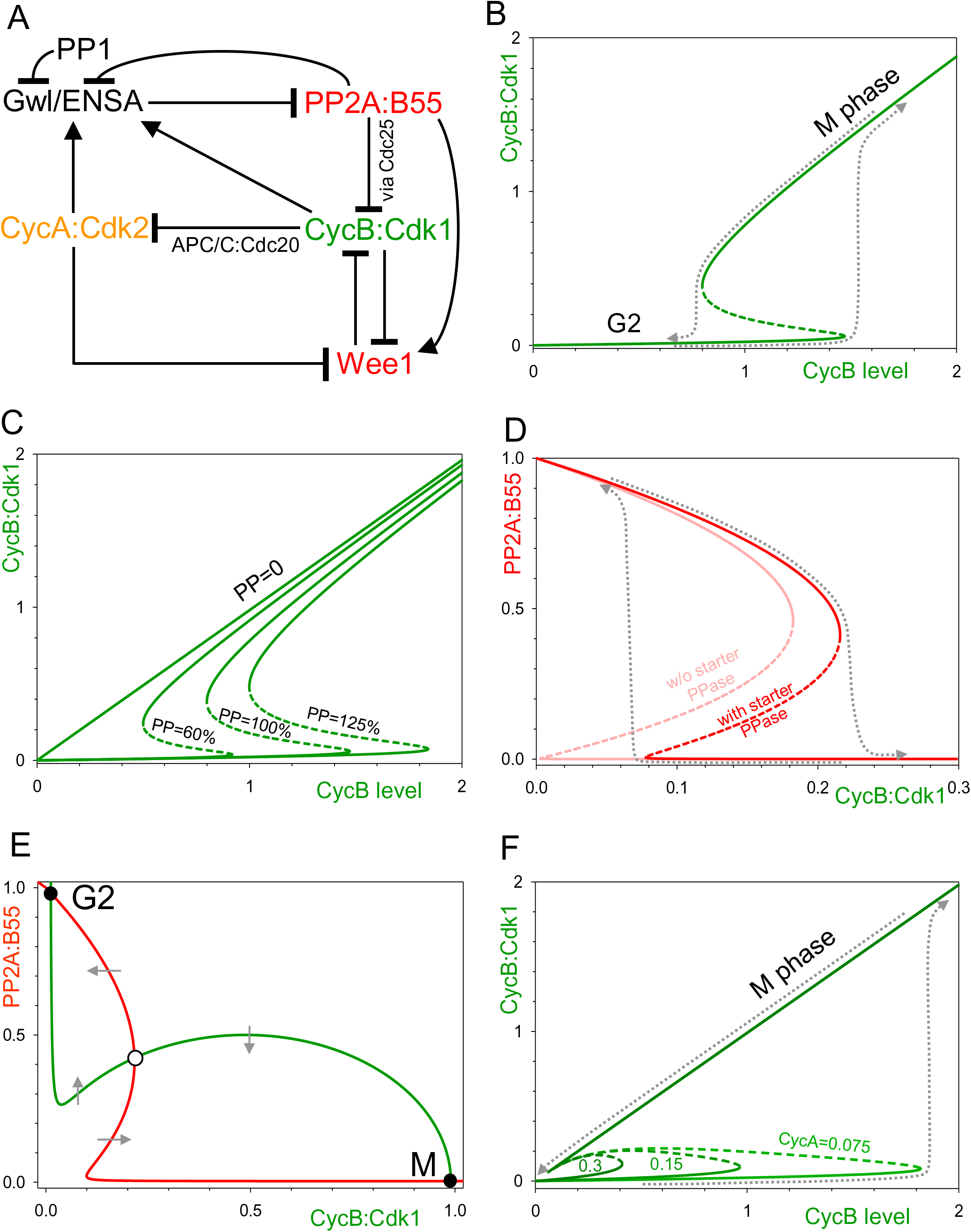
G2/M transition. (A) Influence diagram of the network controlling mitotic entry. The double-negative loop between CycB:Cdk1 and Wee1 is displayed in the diagram, but the double-positive loop between CycB:Cdk1 and Cdc25 is suppressed (for simplicity). The negative influence of CycB:Cdk1 on CycA:Cdk2 is mediated by APC/C:Cdc20. PP2A:B55 has a negative influence on CycB:Cdk1 because PP2A dephosphorylates both Wee1 and Cdc25. The positive influences (the barbed arrows) reflect phosphorylation of Gwl kinase by CycA:Cdk2 and CycB:Cdk1. (B) Signal-response curve (SRC) for CycB:Cdk1 activity (response) as a function of total CycB level (signal), with constant phosphatase (PP) action on Wee1 and Cdc25. The dotted curves indicate the trajectories of the transitions. (C) Rightward shift of the CycB:Cdk1 SRC as a result of increasing phosphatase activity on Wee1 and Cdc25. The larger is the phosphatase activity, the higher is the CycB threshold for CycB:Cdk1 activation. If the phosphatase is inhibited (PP=0), the two thresholds merge, bistability disappears, and Cdk1 activity becomes proportional to CycB level. (D) SRC for PP2A:B55 (response) as a function of CycB:Cdk1 (signal). PP2A:B55 is inactivated at the higher Cdk1 threshold (at mitotic entry) and activated at the lower threshold (at mitotic exit). The starter phosphatase on Gwl kinase increases both thresholds. (E) Intertwined Cdk1 and PP2A:B55 bistable switches create two qualitatively different steady states for interphase and M phase. The PP2A balance curve (red) in this panel is just the PP2A SRC in panel D. The Cdk1 balance curve (green) is a bistable SRC for Cdk1 activity as a function of PP2A activity because of the double-negative feedback loop between Cdk1 and Wee1 (high activity of PP2A pushes this switch into the Wee1-active, Cdk1-inactive steady state). These two balance curves intersect in three steady states: in the interphase steady state (G2), PP2A:B55 is active and CycB:Cdk1 activity is low, while in the mitotic steady state (M) the opposite is true. As before, the grey arrows indicate the directions of the vector field governing the bistable switch. (F) Leftward shift of the CycB:Cdk1 SRC as a result of increasing CycA concentration. See Supplementary Data S2 for details of calculations.

Phosphorylation of Cdk1 substrates remains switch-like in *Xenopus* extracts even if the bistable switch of CycB:Cdk1 activity is disabled by inhibiting Wee1, a fact that highlights the role of Cdk1-counteracting phosphatases [7]. The potential phosphatase candidates are PP1 and PP2A:B55, which are both inhibited by CycB:Cdk1 during mitosis. In particular, phosphorylated endosulfine (P-ENSA) is a strong inhibitory substrate of PP2A:B55, ENSA is phosphorylated by Greatwall (Gwl) kinase, and Gwl is activated by phosphorylation by CycB:Cdk1 during mitosis [8-10]. *In vitro* biochemical reconstitution of the Gwl-ENSA pathway showed that PP2A:B55 counteracts Gwl-activation by Cdk1 and the mutual antagonism between Gwl and PP2A:B55 makes the phosphatase regulation bistable [7] (Figure 2D). The strong inhibition of PP2A:B55 by P-ENSA creates a bootstrapping problem by keeping the phosphatase activation threshold close to zero activity of CycB:Cdk1 (Figure 2D). Timely *in vivo* activation of PP2A:B55 requires the activity of PP1 as Gwl-starter phosphatase [11], thereby shifting the thresholds of the Z-shaped curve to higher Cdk1 values (Figure 2D).

M phase is characterized by extensive phosphorylation of Cdk1 substrates. Cells enter M phase by activating CycB:Cdk1 and inactivating PP2A:B55, and they exit M phase by the reverse transitions. In this scenario, entry into and exit from M phase are controlled by two bistable switches. The kinase and the phosphatase switches are intimately intertwined because PP2A:B55 dephosphorylates not only Gwl but also Wee1 and Cdc25 [10]. Therefore, Cdk1 and the PP2A:B55 mutually inhibit each other’s bistable switching mechanisms, which establishes inverse activities of the kinase and the phosphatase (Figure 2E) and creates amplified changes in the phosphorylation of M phase substrates (low in interphase and high in mitosis).

If indeed mitotic substrate phosphorylation is controlled by bistable switches, then hysteresis should be observed between interphase and mitosis transitions, as has been demonstrated with human cell lines (HeLa and U2OS) using an analogue-sensitive form of Cdk1 (Cdk1as) to control kinase activity [12]. Less Cdk1 inhibitor is required to block the interphase-to-mitosis transition than the opposite transition, supporting the notion of hysteresis. Consistent with the model in Figure 2E, hysteresis of mitotic phosphorylation is only eliminated if both bistable switches are compromised [12].

The CycB threshold of CycB:Cdk1 activation is sensitively dependent on the level of CycA (Figure 2F). In RPE1 cells, the steady state level of CycB in G2 seems to be smaller than the Cdk1-activation threshold in the absence of CycA, which makes CycB:Cdk1 activation fully dependent on CycA [13]. Although CycA helps the G2/M transition, CycA is not required to maintain the mitotic state, since it is well known that CycA is degraded by APC/C:Cdc20 in prometaphase, in a manner that is independent of the mitotic checkpoint. In summary, memory of mitotic entry is provided by these two bistable switches.

### Meta/anaphase transition

The most dramatic cell cycle transition happens at anaphase of mitosis when sister chromatids are irreversibly separated by proteolytic cleavage of cohesin molecules [14]. Cleavage of cohesins is mediated by separase, whose activation requires APC/C:Cdc20, which is controlled by the mitotic checkpoint mechanism. Here, we focus on checkpoint-independent regulation of APC/C:Cdc20 by CycB:Cdk1 (Figure 3A). CycB:Cdk1 phosphorylates both components of the APC/C:Cdc20 complex. Ordered, multi-site phosphorylation of two APC subunits by CycB:Cdk1 delocalizes an inhibitory domain of APC1, which allows Cdc20 to associate to the core APC/C [15,16]. Cdk1 also phosphorylates three N-terminal threonine residues of Cdc20, which inhibits Cdc20 binding to APC/C [17,18]. Since APC/C phosphorylations are activatory, while Cdc20 phosphorylations are inhibitory, CycB:Cdk1 acts on APC/C:Cdc20 complex formation through an **incoherent feedforward loop**. This incoherent feedforward loop is embedded in a **negative feedback loop**, because active APC/C:Cdc20 induces CycB degradation which inactivates Cdk1 (Figure 3A). These regulatory interactions raise the questions of how APC/C:Cdc20 becomes activated at all, and how it maintains its activity until degradation of CycB is complete?

**Figure 3:**
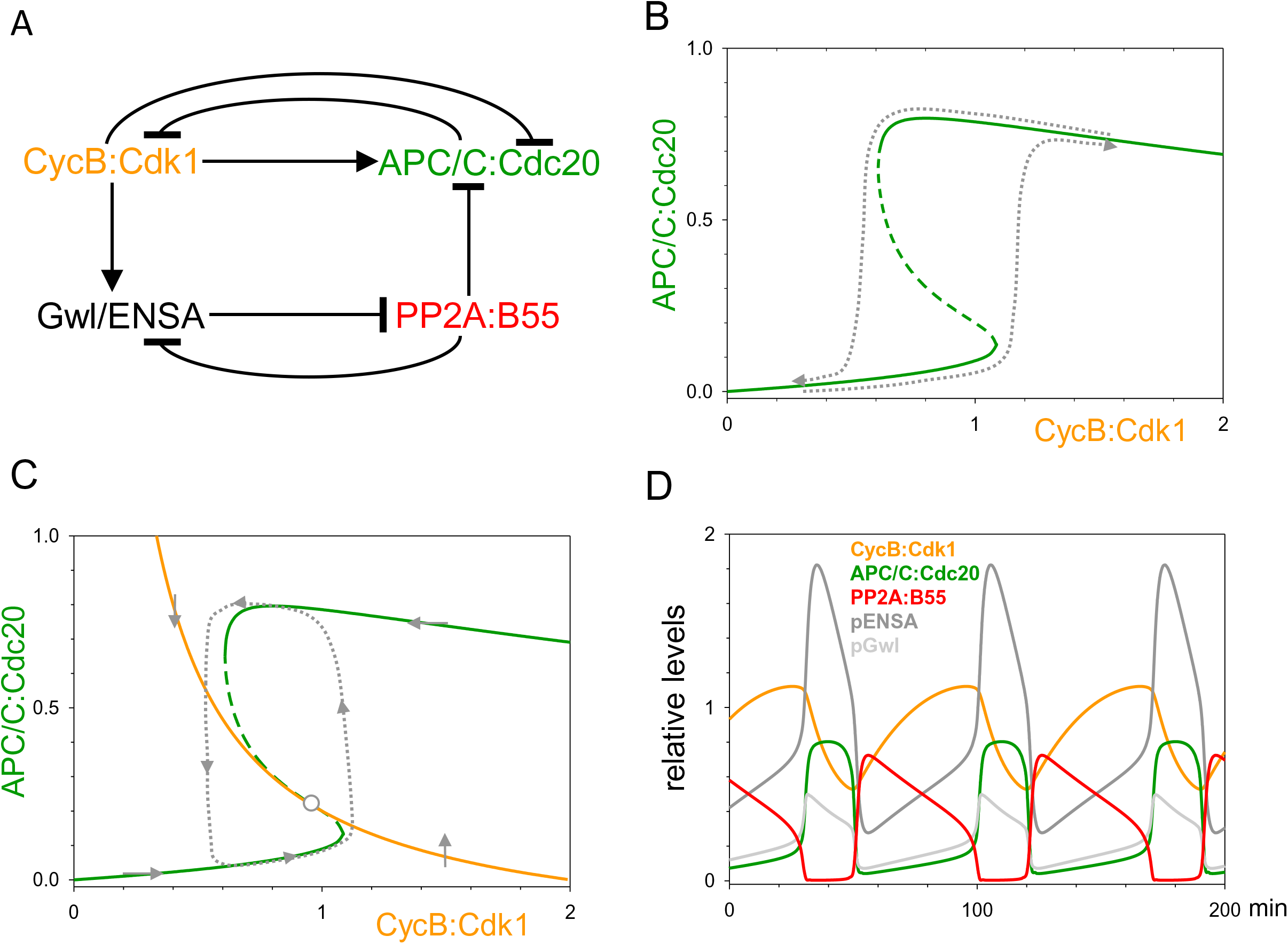
Meta-to-anaphase transition. (A) Influence diagram of the network controlling the meta-to-anaphase transition. (B) Signal-response curve (SRC) for APC/C:Cdc20 (response) as a function of CycB:Cdk1 (signal). The ultrasensitive dependence of APC/C:Cdc20 on CycB:Cdk1 could arise from an underlying toggle switch on PP2A:B55 activity, because PP2A dephosphorylates and inactivates APC/C. Increasing activity of CycB:Cdk1 switches off the activity of PP2A:B55, allowing APC/C:Cdc20 activity to increase abruptly. Bistability predicts that APC/C:Cdc20 is turned off at a smaller CycB:Cdk1 activity. The trajectories of the activation and inactivation transitions are indicated by the dotted curves. (C-D) The interplay between the APC/C:Cdc20 toggle switch and APC/C:Cdc20-induced degradation of CycB creates a positive- and negative-feedback coupled oscillator, as revealed by the balance curves (C) and by temporal simulations (D). The dotted closed curve on (C) indicates the limit cycle oscillation. See Supplementary Data S3 for details of calculations.

In *Xenopus* cell free extracts, where the mitotic checkpoint is not operating, APC/C:Cdc20 activation by CycB:Cdk1 is characterized by a very sharp (ultrasensitive) switch with a characteristic time-delay [19]. This suggests that activatory phosphorylations of APC/C are dominant over Cdc20 inhibitory phosphorylations, and once APC/C:Cdc20 is activated, dephosphorylation of APC/C happens on a slower time-scale than that of Cdc20. The difference in time-scales can be explained by the involvement of different phosphatases or by a preference for Cdc20-P over APC/C-P by the same phosphatase [20,21]. Interestingly all inhibitory phosphorylation sites in Cdc20 are threonine residues, which are preferred by type 2A phosphatases over the phospho-serine residues [20,22] that characterize APC1 activatory sites [16].

We propose that PP2A:B55 is responsible for APC/C dephosphorylation; in which case, PP2A:B55 corresponds to the ‘Inhibitor’ of our generic motif, with APC/C:Cdc20 as the ‘Activator’ of the meta/ana transition (Figure 3A). Since PP2A:B55 is down-regulated by CycB:Cdk1 through the Gwl-ENSA pathway, the activatory phosphorylation of APC/C is regulated by a **coherent feedforward loop** (Figure 3A). The double-negative feedback between A and I of the generic motif is replaced by the mutual antagonism between Gwl and PP2A:B55, which is known to exhibit bistability. According to this proposal, the dependence of APC/C:Cdc20 activity on Cdk1 is governed by a bistable toggle switch (Figure 3B), which might explain the large Hill-exponent (*n*_H_ = 17) of APC/C:Cdc20 activation observed in a Cdk1 dialing-up experiment [19]. In the absence of dialling-down observations, bistability of APC/C:Cdc20 regulation remains a prediction. The APC/C:Cdc20 toggle switch supplemented with the negative feedback of CycB degradation works as an oscillator (Figure 3C, 3D). This oscillation drives the early division cycles in intact *Xenopus* embryos and in cell-free extracts in which the inhibitory phosphorylation of Cdk1 is compromised [23].

### Restriction Point

Before the Restriction Point (RP), entry of mammalian cells into the cell cycle depends on stimulation by growth factors (GFs); after the RP, further progression through the cell cycle is independent of GF until the cell divides [24-26]. Activation of E2F-dependent transcription underlies the RP. E2F is inhibited by the Retinoblastoma (Rb) protein. *i*.*e*., E2F and Rb are generic A and I, respectively, in Figure 4A. The role of pA is played by CycD:Cdk4/6 [27], which is up-regulated by GF. The double-negative feedback between A and I is completed by E2F-induced production of CycE:Cdk2, which hyper-phosphorylates and inactivates Rb. Passage through RP is further accelerated by E2F auto-activating its own transcription (Figure 4A). The expected bistable characteristic of the RP transition has been elegantly demonstrated by the observation that a ten-fold higher concentration of GF is required for E2F induction compared to E2F maintenance [28,29]. According to classical time-lapse experiments, proliferating cells pass through the RP about halfway into G1 phase [25].

**Figure 4:**
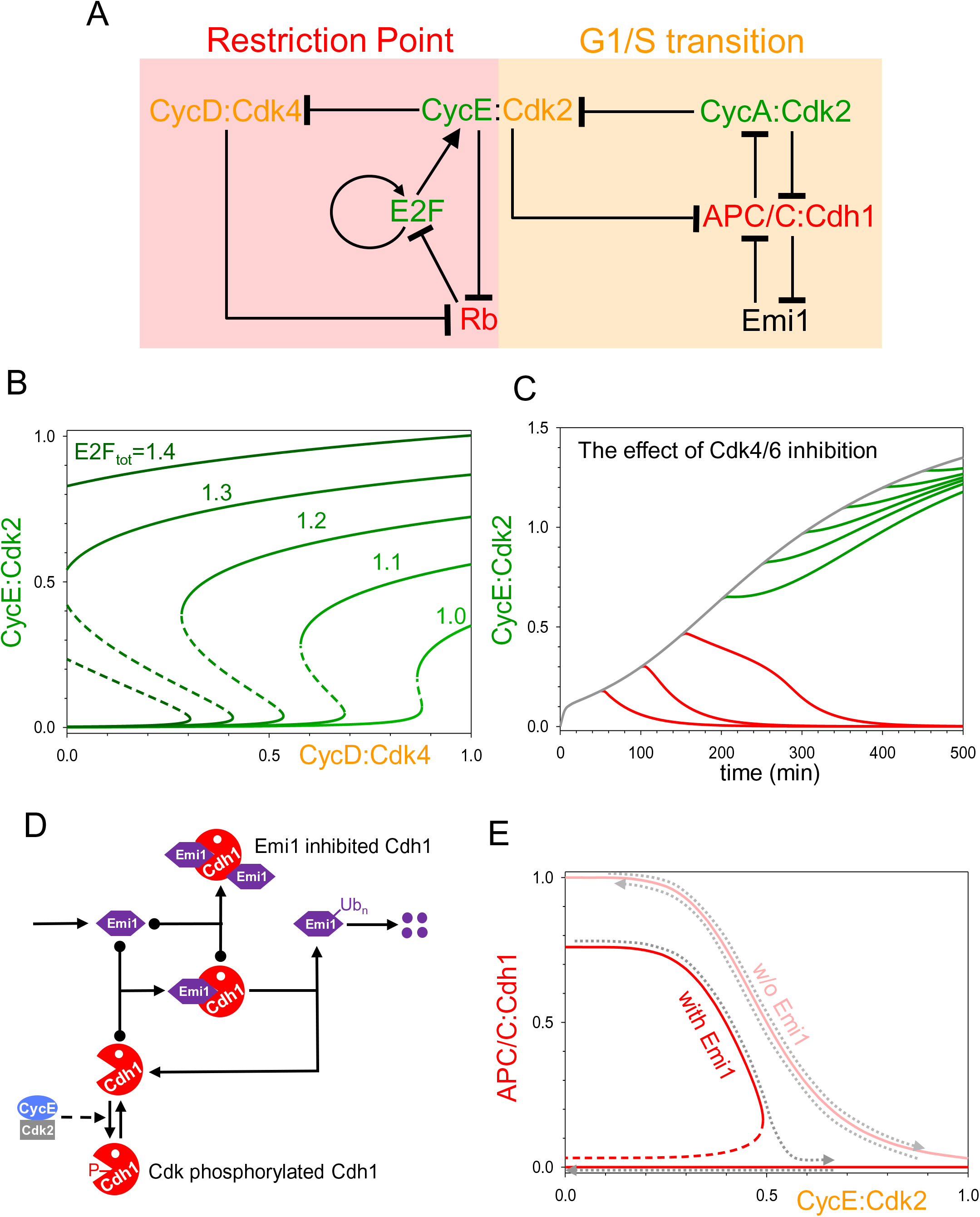
Restriction Point and G1/S transition. (A) Influence diagram of the networks controlling the Restriction Point (RP) and the G1/S transition. (B) Signal-response curves (SRCs) for CycE:Cdk2 (response) as a function of CycD:Cdk4/6 (signal) with different levels of E2F. Notice that increasing level of E2F converts the toggle switch into a one-way switch. (C) Temporal simulation of a bistable Restriction Point model for acute Cdk4/6 inhibition. Dark grey curve: the normal rise in CycE:Cdk2 activity after passing the RP; red and green curves: the effects of acute inhibition of Cdk4/6 introduced at 50 min intervals. The sensitivity of the response (CycE:Cdk2 activity) to Cdk4/6 inhibition (red curves) disappears after 150 min (green curves). (D) Detailed reaction mechanism of APC/C:Cdh1 inhibition at the G1/S transition. (E) SRCs for APC/C:Cdh1 (response) as a function of CycE:Cdk2 activity (signal) in the presence or absence of Emi1. Inhibition of APC/C:Cdh1 by CycE:Cdk2 is ultrasensitive and reversible in the absence of Emi1 (see the parallel running dotted trajectories), but it becomes irreversibly bistable in the presence of Emi1. See Supplementary Data S4 for details of calculations.

Both the cell cycle position of RP and its bistable characteristic have been challenged by recent experiments of Meyer’s group. Using a live-cell sensor for Cdk2 activity, this group has shown that cells bifurcate after cell division into cycling and temporally quiescent states [30]. Immediately after birth, cycling daughter cells start to increase their Cdk2 activity by synthesis of CycE, because they are born with hyper-phosphorylated Rb, indicating that they passed through RP before cell division. In contrast, Cdk2 activity is maintained at low level in Rb hypo-phosphorylated, quiescent cells, which have accumulated p21 (a Cdk inhibitor) due to DNA damage inherited from their mothers [30-33]. The ‘decision’ about whether a daughter cell will be cycling or quiescent cells has been attributed to a ‘calculation’ made by the mother cell, based on competing signals from mitogen-induced synthesis of CycD (pro-cycling) and from DNA damage-induced synthesis of p21 (pro-quiescence) [34].

This bifurcating behaviour has been confirmed for many different human cell lines [35], but it does not seem to be characteristic of primary fibroblasts, which undergo RP during G1 when their Cdk2 activity hits a critical threshold value [36]. It is possible that cell lines growing on culture plates are stimulated by growth factors to attain such a high activity of CycD:Cdk4/6 that Rb does not get dephosphorylated after cell division, unless the cells carry some DNA damage.

The canonical model of the RP proposes that CycD:Cdk4/6 initiates partial Rb phosphorylation, enough to trigger the positive feedback loops whereby CycE:Cdk2 and E2F mutually reinforce each other to induce Rb hyper-phosphorylation. An intuitive consequence of this statement is that Rb phosphorylation should not depend on CycD:Cdk4/6 activity after passage through RP. On the contrary, using acute inhibition of Cdk4/6 by Palbociclib, Meyer’s group showed that Cdk4/6 activity is required to maintain Rb phosphorylation in post-RP daughter cells until they initiate S phase [37]. In other words, CycE/A:Cdk2 activity can only maintain Rb hyper-phosphorylation after the onset of S phase [37].

In our view, these observations do not disqualify the idea of bistability at the RP. If the RP network is a <u>toggle</u> switch, then newborn, post-RP G1 cells are on the upper branch of an S-shaped curve, where E2F is active and CycE is being synthesized (Figure 4B,C). Acute Cdk4/6 inhibition pushes these cells back to the lower branch of the switch, where Rb becomes dephosphorylated and E2F inactivated. Auto-induced E2F synthesis leads to the accumulation of E2F after passing RP, thereby strengthening the feedback loop, as indicated by a decrease of the inactivation threshold (Figure 4B). Increasing E2F level eventually converts the original toggle switch into a one-way switch that is resistant to acute CycD-kinase inhibition (Figure 4B,C).

### G1/S transition

The onset of DNA replication coincides with the appearance of CycA protein in both cancerous and non-cancerous human cell lines [38,39] (Figure 4A), *i*.*e*., CycA:Cdk2 is the generic A of this transition. Although transcription of CycA starts along with CycE after passage through the RP, accumulation of CycA is delayed by APC/C:Cdh1-dependent proteolysis of CycA, because Cdh1 stays active from the end of mitosis until the end of G1 [39,40]. The activity of APC/C:Cdh1 (generic I for this transition) can be inhibited by Cdk1/2-dependent phosphorylation of Cdh1 and by binding to a stoichiometric inhibitor protein, Emi1, whose production is also promoted by E2F. Recent experiments show that Emi1 is not only an inhibitor but also a substrate of APC/C:Cdh1 [40]. Therefore, the initial inactivation of APC/C:Cdh1 relies on CycE:Cdk2 activity (generic pA), which is the only Cdh1 inhibitor among the above mentioned E2F target genes. The initial inactivation of APC/C:Cdh1 leads to partial accumulation of Emi1, and the inhibition of APC/C:Cdh1 by Emi1 allows Emi1 to accumulate further (Figure 4D). Mutual inhibition between APC/C:Cdh1 and Emi1 (APC/C:Cdh1 promotes Emi1 degradation, while high level of Emi1 inhibits the APC/C:Cdh1) has been demonstrated experimentally [40]. This double-negative feedback behaves as a bistable switch, which is flipped to the high Emi1 and low APC/C:Cdh1 side by the rising CycE:Cdk2 activity (Figure 4E). Emi1 is not <u>required</u> for APC/C:Cdh1 inactivation, because CycE:Cdk2 alone is sufficient for inhibiting Cdh1 (Figure 4E). But, in the absence of Emi1 the switch is not bistable, so the Cdk1/2 activity thresholds for turning APC/C:Cdh1 off and on are identical [40]. Emi1 is necessary to make the switch irreversibly bistable (Figure 4E), because in its presence APC/C:Cdh1 cannot be reactivated in post G1/S cells by inhibition of Cdk1/2 [40].

## Conclusions

Recent experimental data summarized above support the crucial role of bistable switches in memory maintenance during cell cycle progression. The history-dependent choice between the two alternative steady states (hysteresis) provides the bistable switch with a memory-storage function. At each cell cycle transition a switch is flipped, which creates a long-lasting, robust, digital-type memory of the executed transition by stabilizing the molecules characteristic of the new cell cycle state.

Our understanding of cell-cycle memory is consistent with and provides further important details to the informal ‘threshold’ model of cell-cycle progression proposed by Stern & Nurse 25 years ago [41]. They suggested that different levels of Cdk1 activity control progression through S phase and mitosis in fission yeast: *‘S phase is initiated when protein kinase activity increases from a very low to a moderate level; maintenance of this moderate level prevents re-initiation of S phase, and a further increase of activity to a high level initiates mitosis*.*’* In our conceptual framework, we may expect a steady increase in cyclin-dependent kinase activity from very low level in G1, to moderate level in G2, to high level in M phase. In addition, our model explains why cells do not easily slip back to an earlier stage of the cell cycle because of the irreversible nature of the bistable switching devices at each transition. Under certain conditions, fission yeast cells can be induced to re-enter S phase without passing through mitosis, as expected for the threshold model [42]. Our mathematical model of bistable cell-cycle transitions in fission yeast provides detailed, quantitative explanations of many different mutant strains exhibiting endoreplication and over-replication of the genome [43].

In addition to their function in memory, bistable switches also play a fundamental role in guaranteeing the correct order of cell cycle events. A necessary requirement of these cell-cycle bistable transitions is that the level of the ‘previous’ activator exceeds the activation threshold of the ‘next’ activator (see Figure 1E-G). Until this condition is satisfied, the bistable switch remains in the stable, pre-transition steady state. Satisfaction of this condition corresponds to a checkpoint for the transition. Cells use this checkpoint property to control progression through the division cycle in response to various situations that may jeopardize proper replication and segregation of chromosomes. A variety of signalling pathways, collectively called ‘surveillance mechanisms,’ monitor diverse threats to genomic integrity and block further progression through the cell cycle until the threat is resolved. Thus, if a previous cell cycle event is not properly completed or conditions are not permissive for the transition, the surveillance mechanism downregulates an activator of the transition, usually by upregulating an inhibitor. In this manner, each cell cycle transition can be checkpoint-arrested if conditions are not satisfactory for further progress. For example, passage through the RP is dependent on mitogen signals in a cell’s environment and is sensitive to DNA damage, which induces a Cdk-inhibitor, p21, as discussed before. The G1/S transition is blocked by DNA damage and in response to other cellular stresses [39]. Complete replication of DNA and proper attachment of all chromosomes to the mitotic spindle control the G2/M and the meta-to-anaphase transitions, respectively. Although the molecular details are specific for each checkpoint, the underlying design principle is the same, namely, blocking the activation of a bistable switch.

## Acknowledgments

We are grateful for profitable discussions with all members of the ‘Bicycle’ group (<u>Bi</u>stability of cell <u>cycle</u> transitions), funded by a BBSRC Strategic LoLa grant (BB/M00354X/1), and especially to Alexis Barr for critically reading the manuscript.

## Notes

### Competing Interest Statement

The authors have declared no competing interest.

