## Supplementary Material for "Mechanisms of signalling-memory governing progression through the eukaryotic cell cycle"

Bela Novak & John J. Tyson

In this Supplementary Material we provide mathematical models that allow the reader to reproduce the Figures in the main text.

#### S1. Generic network motif (Figure 1)

In this simple example, a stoichiometric Inhibitor (I) binds reversibly to an Activator (A) to form an inactive Activator:Inhibitor complex (Comp). A double negative feedback loop is established because the free Activator inactivates the Inhibitor in both its free and complexed form. Inactivation of the Inhibitor is initiated by the previous Activator (pA). The negative feedback by the Activator on the previous Activator is assumed to be nonlinear. The motif can be described dynamically by three ODEs:

$$\frac{dA}{dt} = (k_{dis} + k'_{ai} \cdot [pA] + k_{ai} \cdot [A]) \cdot [Comp] - k_{ass} \cdot ([I]_t - [Comp]) \cdot [A] \quad \text{(S1.1)}$$

$$\frac{d[I]_t}{dt} = k_{aa} \cdot ([I]_{tot} - I_t) - (k'_{ai} \cdot [pA] + k_{ai} \cdot [A]) \cdot [I]_t \quad \text{(S1.2)}$$

$$\frac{d[pA]}{dt} = k_{paa} \cdot ([pA]_{tot} - [pA]) - k_{pai} \cdot [A]^n \cdot [pA] \quad \text{(S1.3)}$$

and the following conservation equations where, $[I]_t$ is the sum of active forms of the Inhibitor (free and complexed with the Activator):

$$[A]_{tot} = [A] + [Comp] = constant$$

$$[I]_{tot} = [I]_{act} + [Comp] + [I]_{inact} = [I]_t + [I]_{inact} = constant$$

Assuming [pA] = constant, we can use Equations (S1.1 &1.2) to calculate the balance curves of Figure 1C, using the nullcline command of XPP-Aut and the following ODE file.

```
# Figure 1C
# Phaseplane for the Activator-Inhibitor double-negative feedback loop
dA/dt = (kdis + kai'*pA + kai*A)*Comp - kass*(It - Comp)*A
dIt/dt = kaa*(Itot - It) - (kai'*pA + kai*A)*It
Comp = Atot - A
p Itot=2, Atot=1, pA=0
p kaa=0.1, kai'=0.1, kai=1, kdis=0.1, kass=100
@ nmesh=400, xp=It, yp=A, xlo=0, ylo=0, xhi=2, yhi=1
done
```

The same equations can be used to calculate the signal-response curve for the Activator (Figure 1D) as a one-parameter bifurcation diagram using XPPAUT with the following ODE file.

```
# Figure 1D
# Signal-response curve for the double-negative feedback loop
dA/dt = (kdis + kai'*pA + kai*A)*Comp - kass*(It - Comp)*A
dIt/dt = kaa*(Itot - It) - (kai'*pA + kai*A)*It
Comp = Atot - A
p pA=0, Itot=2, Atot=1
p kaa=0.1, kai'=0.1, kai=0.5, kdis=0.1, kass=100
@ total=500,dt=1,meth=STIFF,xp=time,yp=A,xlo=0,xhi=500,ylo=0,yhi=1
@ NTST=15,NMAX=100000000,NPR=100000,DS=0.01,BOUNDS=2000
@ DSMAX=0.01,DSMIN=0.001,PARMIN=0,PARMAX=1
@ AUTOXMIN=0,AUTOXMAX=1,AUTOYMIN=0,AUTOYMAX=0.7,AUTOVAR=A
done
```

The phaseplanes on Figure 1E-G can be calculated as nullclines using the following ODE file with pseudo-steady state assumption for the active Inhibitor $[I]_t$.

```
# Figure 1E-G
# Phaseplane for the generic network motif
dpA/dt = kpaa*(pAtot - pA) - kpai*A^n*pA
dA/dt = (kdis + kai'*pA + kai*A)*Comp - kass*(It - Comp)*A
Comp = Atot - A
# Algebraic equation
It = kaa*Itot/(kaa + kai'*pA + kai*A)
# parameter values for the oscillator (Fig.1E)
p Itot=2, Atot=1, pAtot=1
p kaa=0.1, kai'=0.1, kai=0.5, kdis=0.1, kass=100
p kpaa=0.05, kpai=20, n=4
# for the toggle switch use (Fig.1F): pAtot=0.7, kpaa=0.2, kpai=0.5, n=1
# for the one-way switch  use (Fig.1G): kai=0.6
@ nmesh=400, xp=pA, yp=A, xlo=0, ylo=0, xhi=1, yhi=0.8
@ meth=stiff, total=40, dt=0.02
done
```

### S2. G2/M transition and mitotic exit (Figure 2)

The dynamics of CycB:Cdk1 and PP2A:B55 are described by the following ODEs with constant levels of regulatory components (CycB, Gwl, ENSA etc.):

$$\frac{d[Cdk1]}{dt} = V_{25} \cdot ([CycB] - [Cdk1]) - V_{wee} \cdot [Cdk1] \quad \text{(S2.1)}$$

$$\frac{d\,[pGW]}{dt} = (k'_{Cdk} \cdot CycA + k_{Cdk} \cdot [Cdk1]) \cdot ([Gw]_T - [pGW]) - \left(k'_{ppx} + k'_{B55} \cdot [B55]\right) \cdot [pGW] \quad \text{(S2.2)}$$

$$\frac{d\,[pENSA]_t}{dt} = k'_{pGw} \cdot [pGw] \cdot ([ENSA]_{tot} - [pENSA]_t) - k^{B55}_{cat} \cdot ([B55]_{tot} - [B55]) \quad \text{(S2.3)}$$

$$\frac{d[B55]}{dt} = (k_{diss} + k^{B55}_{cat}) \cdot ([B55]_{tot} - [B55]) - k_{ass} \cdot [B55] \cdot \left([pENSA]_t - ([B55]_{tot} - [B55])\right) \quad \text{(S2.4)}$$

and algebraic equations for fast, multisite phosphorylation/dephosphorylation approximated with Hill-functions for Wee1 and Cdc25:

$$V_{wee} = k'_{wee} + (k''_{wee} - k'_{wee}) * \frac{(k_{awee1} \cdot B55)^N}{(k'_{iwee1} \cdot CycA + k_{iwee1} \cdot Cdk1)^N + (k_{awee1} \cdot B55)^N} \quad \text{(S2.5)}$$

$$V_{25} = k'_{25} + \left(k''_{25} - k'_{25}\right) * \frac{(k'_{a25}\cdot CycA + k_{a25}\cdot Cdk1)^M}{(k_{i25}\cdot B55)^M + (k'_{a25}\cdot CycA + k_{a25}\cdot Cdk1)^M} \quad \text{(S2.6)}$$

The easiest way to calculate the signal-response curves for CycB:Cdk1 on Figure 2B,C is by using Equations (S2.1, 2.5 &2.6) and nullcline command of XPP-Aut with the following ODE file.

```
# Signal-response curve for CycB:Cdk1 on Figure 2B & 2C
dCycBT/dt = 0
dCdk1/dt = V25*(CycBT - Cdk1) - Vwee*Cdk1
# Algebraic equations
Vwee   = kwee' + (kwee''-kwee')*(kawee1*PP2AB55)^N/((kiwee1'*CycA + kiwee1*Cdk1)^N + (kawee1*PP2AB55)^N)
V25    = k25' + (k25''-k25')*(ka25'*CycA + ka25*Cdk1)^M/((ki25*PP2AB55)^M + (ka25'*CycA + ka25*Cdk1)^M)
# use PP2AB55 values 0, 0.4, 0.5 and 0.25 for Figure 2C
p PP2AB55=0.4, CycA=0
p k25'=0.02, k25''=1, kwee'=0.02, kwee''=1
p ka25'=1, ka25=1, kiwee1'=1, kiwee1=1
p ki25=1, kawee1=1, kB55'=2.5, N=2, M=2
@ xp=CycBT, yp=Cdk1,xlo=0,xhi=2,ylo=0,yhi=2, nmesh=400
done
```

The following ODE file uses Equations (S2.4) and pseudo-steady state assumptions for pGwl and $pENSA_t$ to calculate the signal-response curves for PP2A:B55 on Figure 2D with the nullcline command of XPP-Aut.

```
# Signal-response curve for PP2A:B55 on Figure 2D
dCdk1/dt = 0
dPP2AB55/dt = (kdiss + kcatB55)*(B55tot - PP2AB55) - kass*PP2AB55*(pENSAt - (B55tot - PP2AB55))
# Algebraic equations
Gwlp = (kCdk'*CycA + kCdk*Cdk1)*Gwtot/(kCdk'*(CycA + Cdk1) + kppx' + kB55'*PP2AB55)
pENSAt = ENSAtot - kcatB55*(B55tot - PP2AB55)/(kpGw'*Gwlp)
p CycA=0, ENSAtot=4, B55tot=1
p kass=3600, kdiss=0.4, kcatB55=3
# use kppx'= 0 & 0.2 to create the two curves
p kpGw'=6, kppx'=0, kCdk'=0.1, kCdk=0.5, Gwtot=1, kB55'=2.5
@ xp=Cdk1, yp=PP2AB55,xlo=0,xhi=0.3,ylo=0,yhi=1, nmesh=400
done
```

The balance curves for PP2A:B55 and CycB:Cdk1 on Figure 2E can be calculated with the following ODE file using Equations (S2.1, 2.4, 2.5 & 2.6) and pseudo-steady state assumptions for pGwl and $pENSA_t$ with the nullcline command of XPP-Aut.

```
# Phaseplane for the G2/M transition on Figure 2E
dCdk1/dt = V25*(CycBT - Cdk1) - Vwee*Cdk1
dPP2AB55/dt = (kdiss + kcatB55)*(B55tot - PP2AB55) - kass*PP2AB55*(pENSAt - (B55tot - PP2AB55))
# Algebraic equations
Gwlp = (kCdk'*CycA + kCdk*Cdk1)*Gwtot/(kCdk'*(CycA + Cdk1) + kppx' + kB55'*PP2AB55)
pENSAt = ENSAtot - kcatB55*(B55tot - PP2AB55)/(kpGw'*Gwlp)
Vwee   = kwee' + (kwee''-kwee')*(kawee1*PP2AB55)^N/((kiwee1'*CycA + kiwee1*Cdk1)^N + (kawee1' +
kawee1*PP2AB55)^N)
V25    = k25' + (k25''-k25')*(ka25'*CycA + ka25*Cdk1)^M/((ki25' + ki25*PP2AB55)^M + (ka25'*CycA + ka25*Cdk1)^M)
p CycA=0, CycBT=1, ENSAtot=4, B55tot=1
p kass=360, kdiss=0.4, kcatB55=3
p kpGw'=6, kppx'=0.2, kCdk'=0.1, kCdk=0.5, Gwtot=1
p k25'=0.01, k25''=1, kwee'=0.01, kwee''=1
p ka25'=1, ka25=1, kiwee1'=1, kiwee1=1
p ki25'=0.1, ki25=1, kawee1'=0.1, kawee1=1, kB55'=2.5, N=2, M=2
@ xp=Cdk1, yp=PP2AB55,xlo=0,xhi=1,ylo=0,yhi=1, nmesh=400
done
```

The effect of CycA on the CycB:Cdk1 signal response curve can be calculated with the following ODE file using Equation S2.1-2.3, 2.5, 2.6 and pseudo-steady state assumptions for the complex between pENSA and PP2A:B55.

```
# The effect of CycA on the CycB:Cdk1 signal-response curve (Figure 2F)
dCdk1/dt = V25*(CycBT - Cdk1) - Vwee*Cdk1
dGwlp/dt = (kCdk'*CycA + kCdk*Cdk1)*(Gwtot-Gwlp) - (kppx' + kB55'*PP2AB55)*Gwlp
dpENSAt/dt = kpGw'*Gwlp*(ENSAtot - pENSAt) - kcatB55*Complex
KM = (kdiss + kcatB55)/kass
BB = pENSAt + B55tot + KM
Complex = 2*pENSAt*B55tot/(BB + sqrt(BB^2 - 4*pENSAt*B55tot))
PP2AB55 = B55tot - Complex
Vwee   = kwee' + (kwee''-kwee')*(kawee1*PP2AB55)^N/((kiwee1'*CycA + kiwee1*Cdk1)^N + (kawee1' +
kawee1*PP2AB55)^N)
V25    = k25' + (k25''-k25')*(ka25'*CycA + ka25*Cdk1)^M/((ki25' + ki25*PP2AB55)^M + (ka25'*CycA + ka25*Cdk1)^M)
# use CycA values of 0.075, 0.15 and 0.3 for the three curves
p CycBT=0, CycA=0, ENSAtot=4, B55tot=1
p kass=360, kdiss=0.4, kcatB55=3
p kpGw'=6, kppx'=0.2, kCdk'=0.1, kCdk=0.5, Gwtot=1
p k25'=0.01, k25''=1, kwee'=0.01, kwee''=1
p ka25'=1, ka25=1, kiwee1'=1, kiwee1=1
p ki25'=0.1, ki25=1, kawee1'=0.1, kawee1=1, kB55'=2.5, N=2, M=2
@ total=500,dt=1,meth=STIFF,xp=time,yp=Cdk1,xlo=0,xhi=500,ylo=0,yhi=1
@ NTST=15,NMAX=100000000,NPR=100000,DS=0.01,BOUNDS=2000
@ DSMAX=0.01,DSMIN=0.001,PARMIN=0,PARMAX=5
@ AUTOXMIN=0,AUTOXMAX=5,AUTOYMIN=0,AUTOYMAX=2,AUTOVAR=Cdk1
done
```

### S3. Meta-to-anaphase transition (Figure 3)

The regulation of APC/C:Cdc20 is described by the following ODEs:

$$\frac{d[Cdk1]}{dt} = k_{scycb} - \left(k'_{dcycb} - k_{dcycb} \cdot [APCpC20]\right) \cdot [Cdk1] \quad \text{(S3.1)}$$

$$\frac{d[APCpC20]}{dt} = k_{a20} \cdot ([APCp] - [APCpC20]) \cdot ([C20] - [APCpC20]) - k_{d20} \cdot [APCpC20] \quad \text{(S3.2)}$$

$$\frac{dAPCp}{dt} = k_{CdkAPC} \cdot [Cdk1] \cdot ([APC]_{tot} - [APCp]) - k_{B55APC} \cdot [B55] \cdot [APCp] \quad \text{(S3.3)}$$

$$\frac{d[C20]}{dt} = k_{dpc20} \cdot ([C20]_{tot} - [C20]) - k_{pc20} \cdot [Cdk1] \cdot [C20] \quad \text{(S3.4)}$$

$$\frac{d\,[pGW]}{dt} = (k'_{Cdk} \cdot CycA + k_{Cdk} \cdot [Cdk1]) \cdot ([Gw]_T - [pGW]) - \left(k'_{ppx} + k'_{B55} \cdot [B55]\right) \cdot [pGW] \quad \text{(S3.5)}$$

$$\frac{d\,[pENSA]_t}{dt} = k'_{pGw} \cdot [pGw] \cdot ([ENSA]_{tot} - [pENSA]_t) - k_{cat}^{B55} \cdot [Complex] \quad \text{(S3.6)}$$

and algebraic equations assuming pseudo-steady state for the complex between PP2A:B55 and ENSA:

$$[Complex] = \frac{2 \cdot [pENSA]_t \cdot [B55]_{tot}}{[pENSA]_t + [B55]_{tot} + K_M + \sqrt{([pENSA]_t + [B55]_{tot} + K_M)^2 - 4 * [pENSA]_t \cdot [B55]_{tot})}} \quad \text{(S3.7)}$$

$$[B55] = [B55]_{tot} - [Complex] \quad \text{(S3.8)}$$

The signal-response curve for APC/C:Cdc20 on Figure 3C & 3D can be calculated as a one-parameter bifurcation diagram using Equations (S3.2-3.8) with the following ODE file.

```
# Signal-response curve for APC/C:Cdc20 on Figure 3C & 3D
dAPCpC20/dt = ka20*(APCp-APCpC20)*(C20-APCpC20) - kd20*APCpC20
dAPCp/dt = kCdkAPC*Cdk1*(APCtot - APCp) - kB55APC*PP2AB55*APCp
dpENSAt/dt = kGwENSA*pGwl*(ENSAtot - pENSAt) - kcatB55*Complex
dpGwl/dt = kCdkGwl*Cdk1*(Gwtot - pGwl) - (kppx' + kB55Gwl*PP2AB55)*pGwl
dC20/dt = kdpc20*(C20tot - C20) - kpc20*Cdk1*C20
BB = pENSAt + B55tot + KM
Complex = 2*pENSAt*B55tot/(BB + sqrt(BB^2 - 4*pENSAt*B55tot))
KM = (kdiss + kcatB55)/kass
PP2AB55 = B55tot - Complex
p Cdk1=0, ENSAtot=4, B55tot=1
p kass=500, kdiss=0.3, kcatB55=1
p kGwENSA=1, kppx'=2
p kCdkGwl=2, kB55Gwl=20, Gwtot=1
p APCtot=1, kCdkAPC=1, kB55APC=20
p C20tot=1, kpc20=1, kdpc20=5, ka20=10, kd20=0.1
@ total=500,dt=0.5,meth=STIFF,xp=time,yp=APCpC20,xlo=0,xhi=500,ylo=0,yhi=1
@ NTST=15,NMAX=100000000,NPR=100000,DS=0.01,BOUNDS=2000
@ DSMAX=0.1,DSMIN=0.001,PARMIN=0,PARMAX=4
@ AUTOXMIN=0,AUTOXMAX=2,AUTOYMIN=0,AUTOYMAX=1,AUTOVAR=APCpC20
done
```

The signal-response curve for Cdk1 on Figure 3D is calculated by the steady state solution of Eq. (S3.1):

$$[Cdk1] = \frac{0.04}{0.02 + 0.1 \cdot [APCpC20]}$$

The trajectory on Figure 3D and the time-course simulation on Figure 3E can be reproduced by the following ODE file.

```
# Temporal oscillation of CycB:Cdk1 and APC/C:Cdc20 on Figure 3E
dCdk1/dt = kscycb - (kdcycb' + kdcycb*APCpC20)*Cdk1
dAPCpC20/dt = ka20*(APCp-APCpC20)*(C20-APCpC20) - kd20*APCpC20
dAPCp/dt = kCdkAPC*Cdk1*(APCtot - APCp) - kB55APC*PP2AB55*APCp
dpENSAt/dt = kGwENSA*pGwl*(ENSAtot - pENSAt) - kcatB55*Complex
dpGwl/dt = kCdkGwl*Cdk1*(Gwtot - pGwl) - (kppx' + kB55Gwl*PP2AB55)*pGwl
dC20/dt = kdpc20*(C20tot - C20) - kpc20*Cdk1*C20
BB = pENSAt + B55tot + KM
Complex = 2*pENSAt*B55tot/(BB + sqrt(BB^2 - 4*pENSAt*B55tot))
KM = (kdiss + kcatB55)/kass
PP2AB55 = B55tot - Complex
aux PP2AB55 = B55tot - Complex
init Cdk1=0.932,APCpC20=0.073,APCp=0.074,pENSAt=0.42,pGwl=0.12,C20=0.84
p ENSAtot=4, B55tot=1
p kass=500, kdiss=0.3, kcatB55=1
p kGwENSA=1, kppx'=2
p kCdkGwl=2, kB55Gwl=20, Gwtot=1
p APCtot=1, kCdkAPC=1, kB55APC=20
p C20tot=1, kpc20=1, kdpc20=5, ka20=10, kd20=0.1
p kscycb=0.04, kdcycb'=0.02, kdcycb=0.1
@ total=200,dt=0.25,meth=STIFF,xp=time,xlo=0,xhi=200,ylo=0,yhi=2
@ NPLOT=5,yp=Cdk1,yp2=APCpC20,yp3=pENSAt,yp4=pGwl,yp5=PP2AB55
done
```

### S4. Restriction Point and G1/S transition (Figure 4)

$$\frac{d[E2F]}{dt} = (k_{dis} + k'_{ai} \cdot [CycD] + k_{ai} \cdot [CycE]) \cdot [Comp] - k_{ass} \cdot ([Rb]_t - [Comp]) \cdot [E2F] - k_{de2f} \cdot [E2F] \qquad \text{(S4.1)}$$

$$\frac{d[Rb]_t}{dt} = k_{aa} \cdot ([Rb]_{tot} - [Rb]_t) - (k'_{ai} \cdot [CycD] + k_{ai} \cdot [CycE]) \cdot [Rb]_t \qquad \text{(S4.2)}$$

$$\frac{d[CycE]}{dt} = k_{se} \cdot [E2F] - k_{de} \cdot [CycE] \qquad \text{(S4.3)}$$

$$\frac{d[E2F]_{tot}}{dt} = k'_{se2f} + k_{se2f} \cdot \frac{E2F^n}{\varepsilon^n + E2F^n} - k_{de2f} \cdot [E2F]_{tot} \qquad \text{(S4.4)}$$

where $[Rb]_{tot} = [RbP] + [Rb] + [Comp]$=$[RbP] + [Rb]_t$ and $[E2F]_{tot} = [E2F] + [Comp]$.

The curves on Figure 4B can be calculated as one-parameter bifurcation diagram with the following ODE file.

```
# Signal-response curves for CycE:Cdk2 for different E2F levels on Figure 4B
dCycE/dt = kse*E2F - kde*CycE
dE2F/dt = (kdis + kai'*CycD + kai*CycE)*Comp - kass*(Rbt - Comp)*E2F
dRbt/dt = kaa*(Rbtot - Rbt) - (kai'*CycD + kai*CycE)*Rbt
Comp = E2Ftot - E2F
# use values of E2Ftot=1 - 1.4 with increments of 0.1
p CycD=0, E2Ftot=1, Rbtot=2
p kaa=0.1, kai'=0.1, kai=0.3, kdis=0.1, kass=100
p kse=0.025, kde=0.025
@ Method=stiff, Total=400, Bounds=100, Dt=1
@ Xplot=t, YPlot=CycE, Xlo=0, Xhi=400, Ylo=0, Yhi=2
@ NTST=150,NMAX=1000000,NPR=10000,DS=0.01,BOUNDS=2000
@ DSMAX=0.01,DSMIN=0.001,PARMIN=-10,PARMAX=1,AUTOVAR=CycE
@ AUTOXMIN=0,AUTOXMAX=1,AUTOYMIN=0,AUTOYMAX=1
done
```

The simulations on Figure 4C can be reproduced by the following ODE file.

```
# Time-course simulations of Cdk4/6 inhibition on Figure 4C
dCycD/dt = -kdcycd*CycD
dE2F/dt = (kdis + kai'*CycD + kai*CycE)*Comp - kass*(Rbt - Comp)*E2F - kde2f*E2F
dRbt/dt = kaa*(Rbtot - Rbt) - (kai'*CycD + kai*CycE)*Rbt
dCycE/dt = kse*E2F - kde*CycE
dE2Ftot/dt = kse2f' + kse2f*E2F^n/(eps^n + E2F^n) - kde2f*E2Ftot
dclock/dt = 1
dkdcycd/dt = 0
Comp = E2Ftot - E2F
global 1 {clock-drug} {kdcycd=alpha}
init CycD=1, E2F=1, Rbt=0, CycE=0, E2Ftot=1, clock=0, kdcycd=0
# the value of 'drug' (0<drug<500) corresponds to the time of inhibitor addition
p drug=1000, Rbtot=2, alpha=1
p kaa=0.1, kai'=0.1, kai=0.3, kdis=0.1, kass=100
p kse=0.025, kde=0.025
p kse2f'=0.005, kse2f=0.005, kde2f=0.005, eps=0.5, n=2
@ nmesh=400, xp=time, yp=CycE, xlo=0, ylo=0, xhi=500, yhi=1.4
@ meth=stiff, total=500, dt=5, bounds=1000
done
```

Emi1 is synthesized and degraded from the complex with Cdh1 (C), unless a second Emi1 is bound (C2). Multisite phosphorylation of Cdh1 by CycE:Cdk2 is described by a power function of CycE:Cdk2 activity. $Cdh1_t$ represents the sum of all unphosphorylated forms (free Cdh1 + C +C2). The G1/S transition is described by the following ODEs:

$$\frac{d[Emi1]_{tot}}{dt} = k_{semi} - k'_{demi} \cdot [Emi1]_{tot} - k_{demi} \cdot [C] \quad \text{(S4.5)}$$

$$\frac{d[Cdh1]_t}{dt} = k_{dp}^n \cdot ([Cdh1]_{tot} - [Cdh1]_t) - k_p \cdot CycE^n \cdot [Cdh1]_t \quad \text{(S4.6)}$$

$$\frac{d[C]}{dt} = k_{ass} \cdot [Cdh1] \cdot [Emi1] - (k_{diss} + k_{ass2} \cdot [Emi1] + k'_{demi} + k_{demi} + k_p \cdot CycE^n) \cdot [C] + k_{diss2} \cdot [C2] \quad \text{(S4.7)}$$

$$\frac{d[Cdh1]}{dt} = (k_{diss} + k'_{demi} + k_{demi}) \cdot [C] + k'_{demi} \cdot [C2] + k_{dp}^n \cdot [pCdh1] - k_{ass} \cdot [Cdh1] \cdot [Emi1] - k_p \cdot CycE^n \cdot [Cdh1] \quad \text{(S4.8)}$$

and the following algebraic equations: $[Emi1] = [Emi1]_{tot} - [C] - 2 \cdot [C2]$ and $[C2\,] = [Cdh1]_t - [C] - [Cdh1]$.

Figure 4E can be reproduced by the ODE file below by setting $k_{semi}$ to 0.1 and zero.

```
# Signal-response curve for the one-way switch at G1/S transition (Figure 4D)
dEmi1tot/dt = ksemi - kdemi'*Emi1tot - kdemi*C
dCdh1t/dt = kdp^n*pCdh1 - kp*CycE^n*Cdh1t
dC/dt = kass*Cdh1*(Emi1tot-C-2*C2) - (kdiss+kass2*(Emi1tot-C-2*C2)+kdemi'+kdemi+kp*CycE^n)*C + kdiss2*C2
dCdh1/dt = (kdiss+kdemi'+kdemi)*C+kdemi'*C2+kdp^n*pCdh1-kass*Cdh1*(Emi1tot-C-2*C2)-kp*CycE^n*Cdh1
# Algebraic equations
pCdh1 =Cdh1tot - Cdh1t
C2 = Cdh1t - C - Cdh1
init Emi1tot=0.288, Cdh1t=0.99, C=0.19, Cdh1=0.76
p CycE=0, kp=1, kdp=0.5, n=5
p ksemi=0.1, kdemi'=0.01, kdemi=0.5
p Cdh1tot=1, kass=100, kdiss=0.5
p kass2=10, kdiss2=0.1
@ total=500,dt=0.5,meth=STIFF,xp=time,yp=Emi1tot, xlo=0,xhi=500,ylo=0,yhi=1
@ NTST=15,NMAX=1000000,NPR=10000,DS=0.01,BOUNDS=2000
@ DSMAX=0.1,DSMIN=0.02,PARMIN=-10,PARMAX=1, AUTOVAR=Cdh1
@ AUTOXMIN=0,AUTOXMAX=1,AUTOYMIN=0,AUTOYMAX=1
done
```
